# Vertical transmission of chronic wasting disease in free-ranging white-tailed deer populations

**DOI:** 10.1101/2025.01.24.634834

**Authors:** Audrey M. Sandoval, Amy V. Nalls, Erin E. McNulty, Nathaniel D. Denkers, Devon J. Trujillo, Zoe Olmstead, Ethan Barton, Jennifer R. Ballard, Daniel M. Grove, Jeremy S. Dennison, Natalie Stilwell, Christopher A. Cleveland, James M. Crum, Mark G. Ruder, Candace K. Mathiason

## Abstract

Chronic wasting disease (CWD) is a fatal neurodegenerative disease affecting cervids across North America, Northern Europe, and Asia. Disease transmission among cervids has historically been attributed to direct animal-to-animal contact with ‘secreta’ (saliva, blood, urine, and feces) containing the infectious agent, and indirect contact with the agent shed to the environment in these bodily components. Mounting evidence provides another mechanism of CWD transmission, that from mother-to-offspring, including during pregnancy (vertical transmission). Here we describe the detection of the infectious CWD agent and prion seeding in fetal and reproductive tissues collected from healthy-appearing free-ranging white-tailed deer (*Odocoileus virginianus*) from multiple U.S. states by mouse bioassay and *in vitro* prion amplification assays. This is the first report of the infectious agent in several *in utero* derived fetal and maternal-fetal reproductive tissues, providing evidence that CWD infections are propagated within gestational fetal tissues of white-tailed deer populations. This work confirms previous experimental and field findings in several cervid species supporting vertical transmission as a mechanism of CWD transmission and helps to further explain the facile dissemination of this disease among captive and free-ranging cervid populations.

## INTRODUCTION

Chronic wasting disease (CWD) is an efficiently transmitted prion disease of captive and free-ranging cervid species (deer, elk, moose, reindeer) that continues to be detected in parts of North America, Scandinavia, and Asia^1-4^. Transmissible spongiform encephalopathies (TSEs), including CWD, affect the central nervous system causing slow and progressive neurological damage resulting in behavioral changes, wasting, and eventual death^5^. It has been confirmed that infectious CWD prions are shed in ‘secreta’ (saliva, blood, urine and feces) during the characteristic long asymptomatic phase of disease, which can range from months to years^6-9^. Prions shed in secreta to the environment and those in the carcass after death have robust stability, withstanding degradation and remaining infectious for many years ^10-13^. Thus, the facile dissemination of CWD among cervids has largely been attributed to contact with the infectious agent via direct animal-to-animal contact (e.g., grooming, breeding, birthing fluids) and indirect interactions with the infectious agent shed to the environment (e.g., foraging, tree rubs, mineral licks, soil consumption)^14-18^. What remains less understood is the role of maternal CWD infections and mother-to-offspring transmission.

Previous studies have reported that offspring born to prion-infected deer, cattle, and sheep are at risk for developing CWD, bovine spongiform encephalopathy (BSE), and scrapie, respectively^19-21^. Maternal transmission of feline spongiform encephalopathy (FSE) has been implicated in a seven-year-old cheetah born to an FSE-infected mother that was raised in a TSE-free environment yet developed FSE^22^. To date there are no reports of human Creutzfeldt-Jakob disease (CJD) mother-to-offspring transmission, yet, prion deposition and infectivity have been demonstrated in human reproductive tissues and cord blood of variant CJD (vCJD) patients^23,24^. Several additional studies provide strong evidence for *in utero* or vertical prion transmission. *In vitro* prion deposition has been detected in reproductive tissues (ovary, uterus, placentomes, and amniotic fluid) of scrapie-infected sheep^25-27^.

We first described CWD exposure prior to parturition upon the demonstration of PrP^CWD^ immunohistochemical (IHC) deposition within tonsillar lymphoid biopsies as early as 41 days post parturition in offspring born to experimental CWD-infected Reeves’ muntjac (*Muntiacus reevesi*) dams^28^. In these studies, amplifiable prions were demonstrated in multiple maternal reproductive tissues/fluids and fetal samples harvested throughout gestation demonstrating that *in utero* transmission could occur at any stage of gestation or CWD infection^28,29^. More importantly, we demonstrated the presence of the infectious prion agent within tissues of the maternal pregnancy microenvironment (placentomes, uterus, amniotic fluids) that had the capacity to initiate CWD infections and disease progression when inoculated into transgenic mice^29^. What remained unknown is if these findings would hold true for naturally infected, free-ranging cervid species that most assuredly receive smaller doses of CWD in their natural environment. Subsequently, we and others have detected amplifiable prions in dam reproductive and fetal tissues harvested from naturally exposed CWD-infected Rocky Mountain elk (*Cervus canadensis nelsoni*) and white-tailed deer (WTD; *Odocoileus virginianus*)^30-32^. What has not been shown is if the prion detection in these tissues is infectious. Thus, an overarching question has been, does prion deposition within gestational tissues of free-ranging cervids result in fetal prion infections?

Here, we assessed maternal reproductive tissues/fluids, and *in utero* derived fetal tissues from free-ranging WTD naturally exposed to CWD for the presence of prion seeding activity (prions) by amplification assays (protein misfolding amplification assay [PMCA] and real-time quaking induced conversion [RT-QuIC]). More importantly, to determine the biological relevance of *in vitro* prion detection, we assessed reproductive and fetal tissues for the presence of infectivity by bioassay in cervid *Prnp* transgenic mice. Bioassay is the only way to assess for the presence of the infectious agent that is known to lead to the initiation and progression of prion infections.

To account for the potential that prion protein (*Prnp*) genotype may impact vertical transmission, as has been demonstrated in sheep scrapie^25^, and is well known to affect cervid prion susceptibility^33,34^, disease progression and shedding^34-36^, we assessed the *Prnp* genotypes at codon 95 & 96 of both dams and fetuses, and corelated these findings to CWD disease status. Finally, as previous experimental studies in the native host demonstrate that CWD high dose saliva, blood and fomite exposures resulted in first CWD positive status between 12 and 19 months post-inoculation^15^, we retrospectively examined historical CWD surveillance data from three study sites in Arkansas, Tennessee, and West Virginia to look for evidence of natural infection in WTD fawns less than 12 months of age using tests of similar sensitivity (western blot and ELISA respectively)^37^.

We report the presence of the infectious CWD agent within the pregnancy microenvironment and fetal tissues of free-ranging, naturally exposed WTD (i.e. vertical transmission). This demonstrates that CWD can cross the maternal-fetal interface to initiate an active gestational infection. We also show that *Prnp* polymorphisms may impact vertical transmission, as there were no cases of fetal positivity when codon 96GS polymorphism was present in the maternal/fetal pairs. Furthermore, examination of historical CWD surveillance data revealed infections in WTD fawns between 6-10 months of age, suggesting an earlier exposure to CWD than postpartum maternal saliva, blood, or environmental contact^15^. These findings lend biological relevance to earlier detection of *in vitro* prion seeding activity within maternal reproductive and fetal tissues of experimental and free-ranging cervids^28,30-32^, and demonstrates that *in utero*/vertical transmission is another factor in the transmission dynamics of CWD in the native cervid host.

## RESULTS

### CWD detected in free-ranging white-tailed deer dams from Arkansas, Tennessee, and West Virginia

To survey the extent of CWD dissemination within free-ranging WTD, we evaluated peripheral lymphoid tissues (RPLN, tonsil, RAMALT, and third eyelid) and central nervous system tissue (obex) from a total of n=31 WTD dams by IHC and RT-QuIC (n=28 from CWD endemic regions and n=3 from a CWD nonendemic region). From our CWD endemic study cohort (n=28), we detected CWD seeding activity and/or PrP^cwd^ deposition in 16 of 28 (57.1%) RPLN, 14 of 28 (50%) tonsil, 12 of 28 (42.8%) RAMALT, 11 of 28 (39.2%) third eyelid, and 8 of 28 (28.6%) obex tissues (**Table 1**). The detection of CWD by RT-QuIC amyloid seeding preceded or coincided with IHC detection in all but two samples (**Table 1**). Additionally, in dams with sufficient secreta collected to assess prion shedding (n=27 saliva; n=17 urine), amyloid seeding was detected in 3 of 27 (11%) saliva samples and 0 of 17 (0%) urine samples (**Table 1**). Overall, 16 of the 28 dams were CWD-positive on one or more tests with results from each state as follows: 3 of 10 (30%) Arkansas, 8 of 10 (80%) Tennessee, and 5 of 8 (62.5%) West Virginia. All negative control deer harvested in Georgia (n=3) had no detectable prions in all samples tested.

**Table 1.** Summary of pregnant white-tailed deer (WTD) data including age, genotype based on *Prnp* codon 96, and chronic wasting disease (CWD) status of lymphoid tissues, obex and secreta. RT-QuIC and IHC status of tissues including retropharyngeal lymph node (RPLN), third eyelid, recto-anal mucosal lymphoid tissue (RAMALT), and obex. RT-QuIC status of secreta including urine and saliva. Data are separated by location (state) of sample collection. Samples from WTD in Georgia, an area with no CWD detections, remained negative and served as sample comparison. NA, not applicable – sample was unavailable for testing. Age estimated based on tooth wear and replacement.

| Pregnant Doe |  |  |  |  |  |  |  |  |  |  |  |  |  |  |
| --- | --- | --- | --- | --- | --- | --- | --- | --- | --- | --- | --- | --- | --- | --- |
| Location | Dam ID | Age (years) | Geno-type (96) | RPLN |  | Tonsil |  | RAMALT |  | Third Eyelid |  | Obex |  | Secreta (Urine & Saliva) |
|  |  |  |  | QuIC | IHC | QuIC | IHC | QuIC | IHC | QuIC | IHC | QuIC | IHC |  |
| Georgia | 4-2 | 1.75 | GG | - | - | - | - | - | - | - | - | - | - | - |
|  | 4-3 | 6.75 | GS | - | - | - | - | - | - | - | - | - | - | - |
|  | 4-4 | 3.75 | GG | - | - | - | - | - | - | - | - | - | - | NA |
| Arkansas | 5-1 | 4.75 | SS | - | - | - | - | - | - | - | - | - | - | - |
|  | 5-2 | 2.75 | GG | - | - | - | - | - | - | - | - | - | - | NA |
|  | 5-3 | 3.75 | GG | - | - | - | - | - | - | - | - | - | - | - |
|  | 5-4 | 2.75 | GG | - | - | - | - | - | - | - | - | - | - | - |
|  | 5-6 | 4.75 | GG | + | + | + | + | + | NA | + | NA | + | + | - |
|  | 5-7 | >8.5 | GG | - | - | - | - | - | - | - | - | - | - | NA |
|  | 5-8 | 2.75-3.75 | GG | - | + | - | - | + | - | - | - | - | - | - |
|  | 5-9 | >5.5 | GG | - | - | - | - | - | - | - | - | - | - | - |
|  | 5-10 | 2.34 | GS | - | - | - | - | - | - | - | - | - | - | - |
|  | 5-12 | 2.75 | GG | + | + | + | - | + | - | + | - | + | + | - |
| Tennessee | 6-1 | 2.75 | GG | + | + | + | + | + | + | + | + | + | + | + |
|  | 6-3 | 2.75 | GG | - | - | - | - | - | - | - | - | - | - | - |
|  | 6-5 | 2.75 | GG | + | + | + | + | + | + | + | + | + | + | + |
|  | 6-6 | 3.75 | GG | + | + | + | - | - | - | - | - | - | - | - |
|  | 6-10 | 2.75 | GG | + | + | + | + | + | + | + | NA | + | + | - |
|  | 6-11 | 4.75 | GS | - | - | - | - | - | - | - | - | - | - | - |
|  | 6-12 | >5.75 | GS | + | - | - | - | - | - | - | NA | - | - | - |
|  | 6-13 | 2.75 | GS | + | + | + | + | + | + | + | + | - | - | - |
|  | 6-14 | 2.75 | GG | + | - | + | - | - | - | - | - | - | - | - |
|  | 6-15 | 2.75 | GS | + | - | + | + | + | + | + | - | - | + | + |
| West Virginia | 7-1 | 1.8 | GG | - | - | - | - | - | - | - | - | - | - | NA |
|  | 7-2 | 6.8 | GS | + | + | + | + | + | - | + | - | - | - | - |
|  | 7-3 | 2.8 | GG | + | + | + | + | + | + | + | - | - | - | - |
|  | 7-4 | 1.8 | GG | + | + | + | + | + | + | + | + | + | + | NA |
|  | 7-6 | 3.8 | GG | + | + | + | - | - | - | - | NA | - | - | NA |
|  | 7-7 | 4.8 | GS | - | - | - | - | - | - | - | - | - | - | - |
|  | 7-9 | 2.8 | GS | - | - | - | - | - | - | - | - | - | - | NA |
|  | 7-10 | 2.8 | GG | + | + | + | + | + | + | + | + | + | + | NA |
| Total Positive |  |  |  | 57% |  | 50% |  | 43% |  | 40% |  | 29% |  | 14% |

### *In vitro* prion seeding present within the pregnancy microenvironment of free-ranging white-tailed deer

We assessed the pregnancy microenvironment of this study cohort, including negative control dams, for the presence of prions using RT-QuIC. We evaluated uterine tissue from all 31 dams, a total of 51 amniotic fluid samples, and 200 placentomes. From the 16 CWD positive dams, prion seeding activity was found in 5 of 16 uterine tissues (31.25%), 5 of 48 amniotic fluid samples (10.4%) and 18 of 116 placentomes (15.5%) (**Figure 1 a-c; Table 2**). Of note, 17 of 18 (94.4%) positive placentomes correlated to dams with positive uterine tissue (**Table 2**). By state, we report positive seeding activity in the reproductive tissues and fluids of dams from Arkansas (2/10; 20%), Tennessee (3/10; 30%), and West Virginia (3/8; 37.5%). Reproductive tissues (uterus, placentome, amniotic fluid) collected from our study cohort dams that were lymphoid and obex RT-QuIC negative (n=12), as well as dams harvested from a CWD non-endemic region (Georgia; n=3) (**Table 1**), remained RT-QuIC negative. Overall, of the 16 confirmed CWD-positive free-ranging dams, *in vitro* amyloid seeding was demonstrated in the pregnancy microenvironment of 8 of 16 (50%).

**Table 2.**
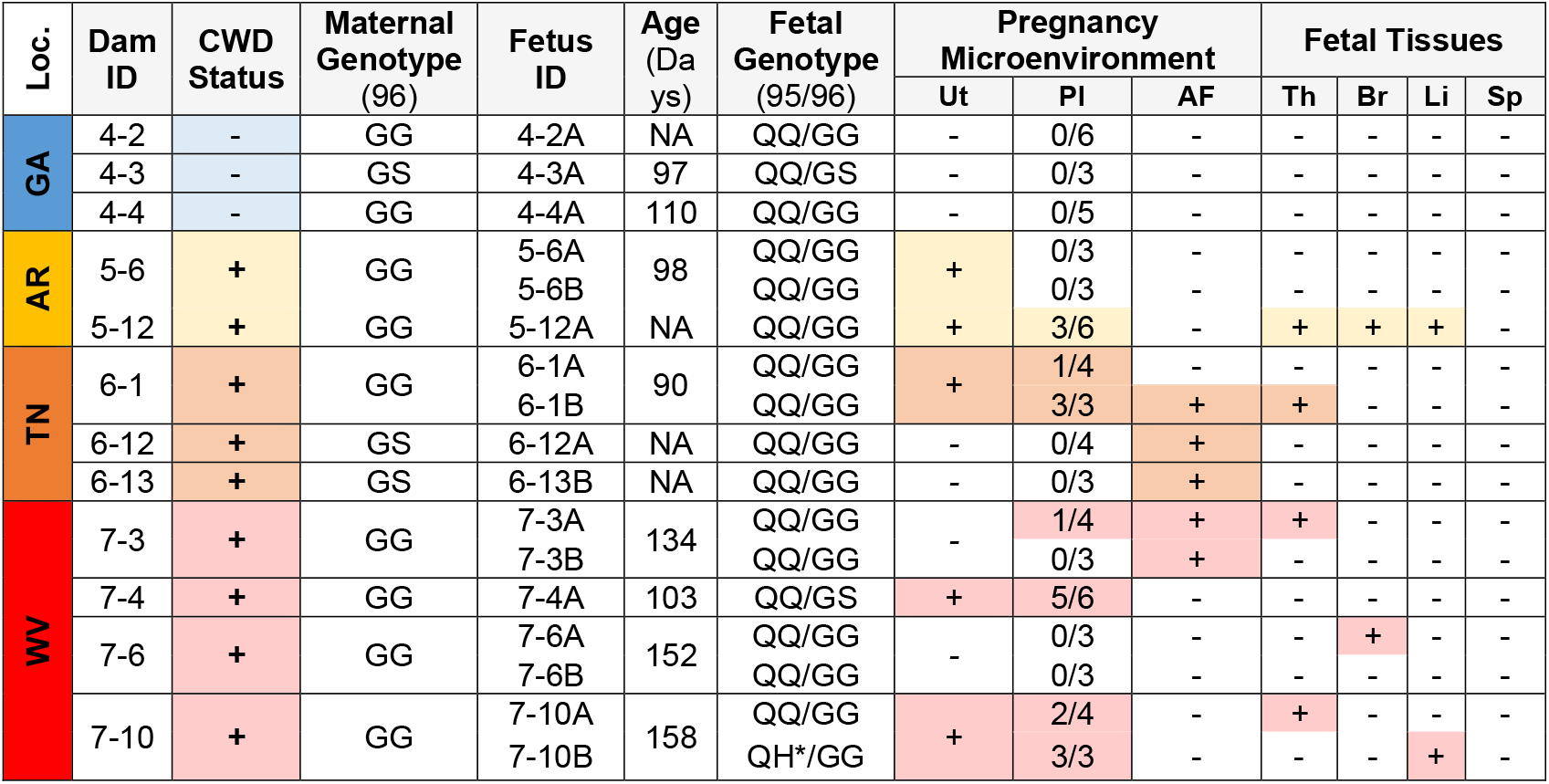
Condensed data from CWD-positive, female white-tailed deer, showing prion seeding activity in their pregnancy microenvironment and/or fetal tissues based on RT-QuIC. CWD status is based on one or more tissue samples testing positive by RT-QuIC or IHC, as shown in Table 1. Results from deer that tested negative by RT-QuIC in their pregnancy microenvironment and fetal tissues are available in supplementary data. Data are separated by location (state) of sample collection, including Georgia (GA), blue; Arkansas (AR), yellow; Tennessee (TN), orange; West Virginia (WV), red. Abbreviations as follows: (Ut)erus, (Pl)acentome, (A)mniotic (F)luid, (Th)ymus, (Br)ain, (Li)ver, & (Sp)leen. Placentome data shown as #pos/total placentomes. Gestational age estimated by Hamilton equation. NA, not applicable – data point not collected during necropsy. Genotype representative of *Prnp* 96. * 7-10B fetus was QH heterozygous at codon 95, all other fetuses, except heterozygous 5-9B (not shown), were QQ at the same position.

**Figure 1.**
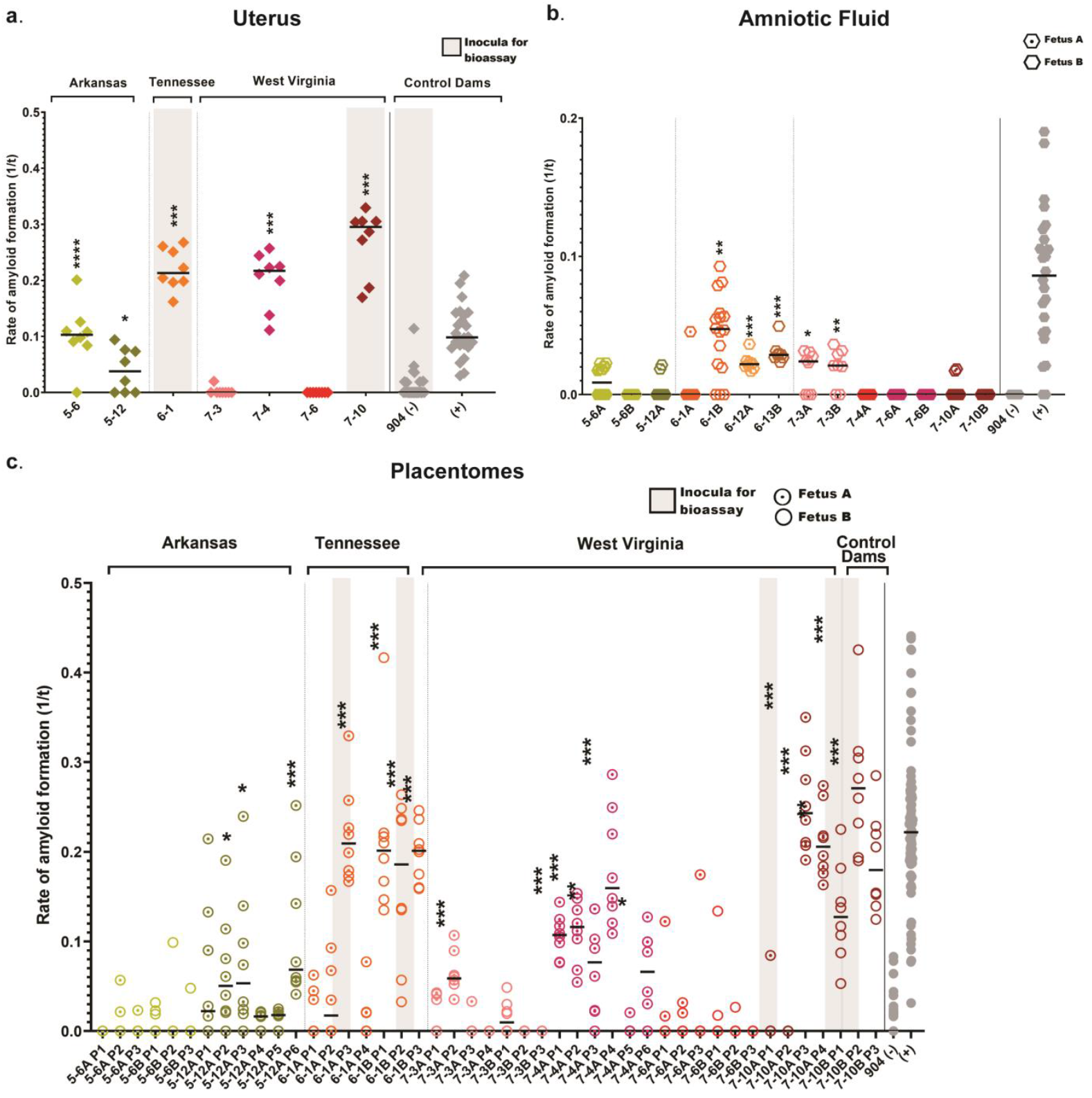
CWD detection in female white-tailed deer in the pregnancy microenvironment determined by RT-QuIC. Grouped based on state of collection. Amyloid seeding in one or more of the following tissues: uterus, amniotic fluid, and placentomes (a-c; Table 2) from CWD-positive dams as determined by RT-QuIC & IHC (Table 1). Amyloid formation displayed as reaction rates (1/time (t) to ThT fluorescent threshold). Horizontal lines represent the medians with n=8 replicates (dam tissues). Negative controls from all Georgia deer (n=3) remained negative in all samples tested and CWD-positive controls sourced from experimentally inoculated muntjac (last data point on the right). P values as follows: * <0.05, ** <0.01, *** <0.001, ****<0.0001. Grey shaded rectangles in panels A and C denote samples utilized for bioassay inoculations (Table 3).

### *In vitro* prion seeding present in fetal tissues harvested from free-ranging white-tailed deer

After demonstrating positive seeding activity in tissues of the maternal reproductive tract (uterus) and at the maternal-fetal interface (placentomes and amniotic fluid), we evaluated 54 fetuses (CWD endemic (n=51); nonendemic (n=3)) with sufficient tissue available [thymus (n=54), brain (n=54), liver (n=49), spleen (n=49)] for the presence of CWD by PMCA^38^. Given the small size of each fetus and thus limited sample volume harvested, in conjunction with presumed low CWD burdens within each tissue type^32^, we combined PMCA amplification with RT-QuIC readout^37^ to maximize detection sensitivity. Here we detected PMCA amplifiable amyloid seeds in thymus tissue harvested from four fetuses (4/51; 7.8%), brain tissue harvested from two fetuses (2/51; 3.9%), and liver tissues from two fetuses (2/46; 4.4%) (**Figure 2**). Fetal spleen tissue did not reveal PMCA amplification (**Table 2**). All CWD negative dams carried fetuses that remained negative for PMCA amplification (**Table 1**). We report a unique case in which we found PMCA amplification in fetal brain tissue (**Figure 2**) harvested from a free-ranging CWD-positive dam that did not have evidence of CWD dissemination within the reproductive tissues tested (**Figure 1 a-c; Table 2**). To summarize, 5 of the 16 (31.3%) free-ranging CWD-positive dams were carrying at least one fetus with tissues that harbored PMCA amplifiable prion seeds, with one each from Arkansas and Tennessee, and three from West Virginia.

**Table 3.**
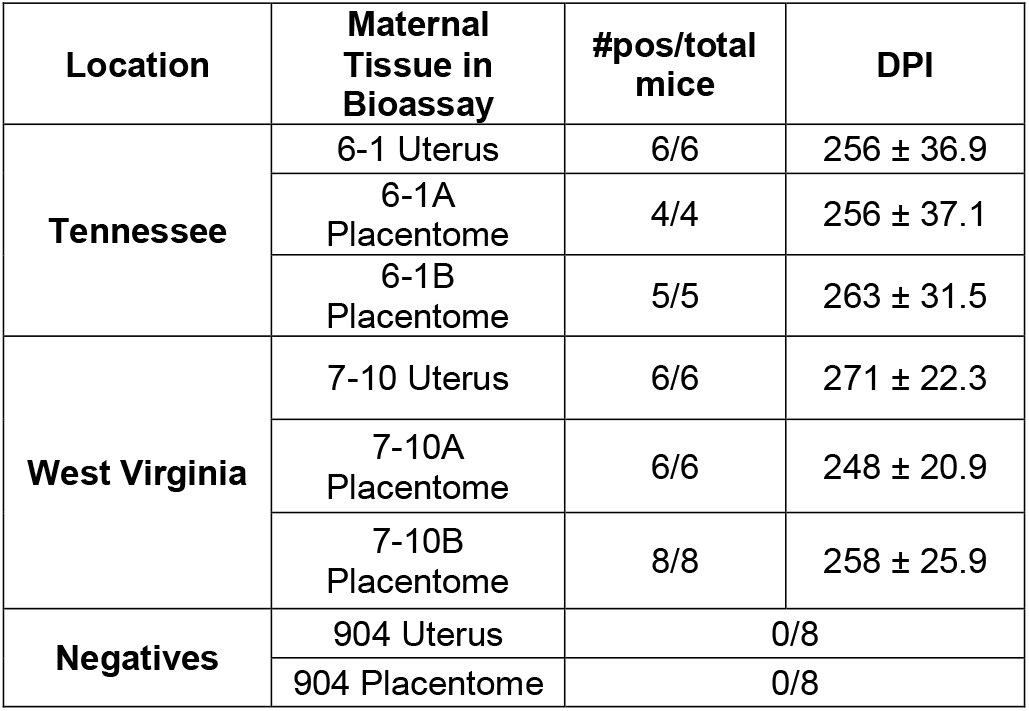
Mouse bioassay results for CWD-positive white-tailed deer maternal reproductive tissues. Cervidized (Tg^CerPrP-E226^5037^+/-^) mice were intracranially (IC) inoculated with uterus (dams 6-1 and 7-10) and placentome (dams 6-1A and B and 7-10A and B) samples. Terminal mouse brains were tested by conventional RT-QuIC for amyloid seeding activity. Mock inoculated mice remained free of clinical signs throughout, and tested CWD-negative upon terminal collections in all cohorts. DPI = days post inoculation. # pos/total mice = number CWD-positive/total number inoculated.

**Figure 2.**
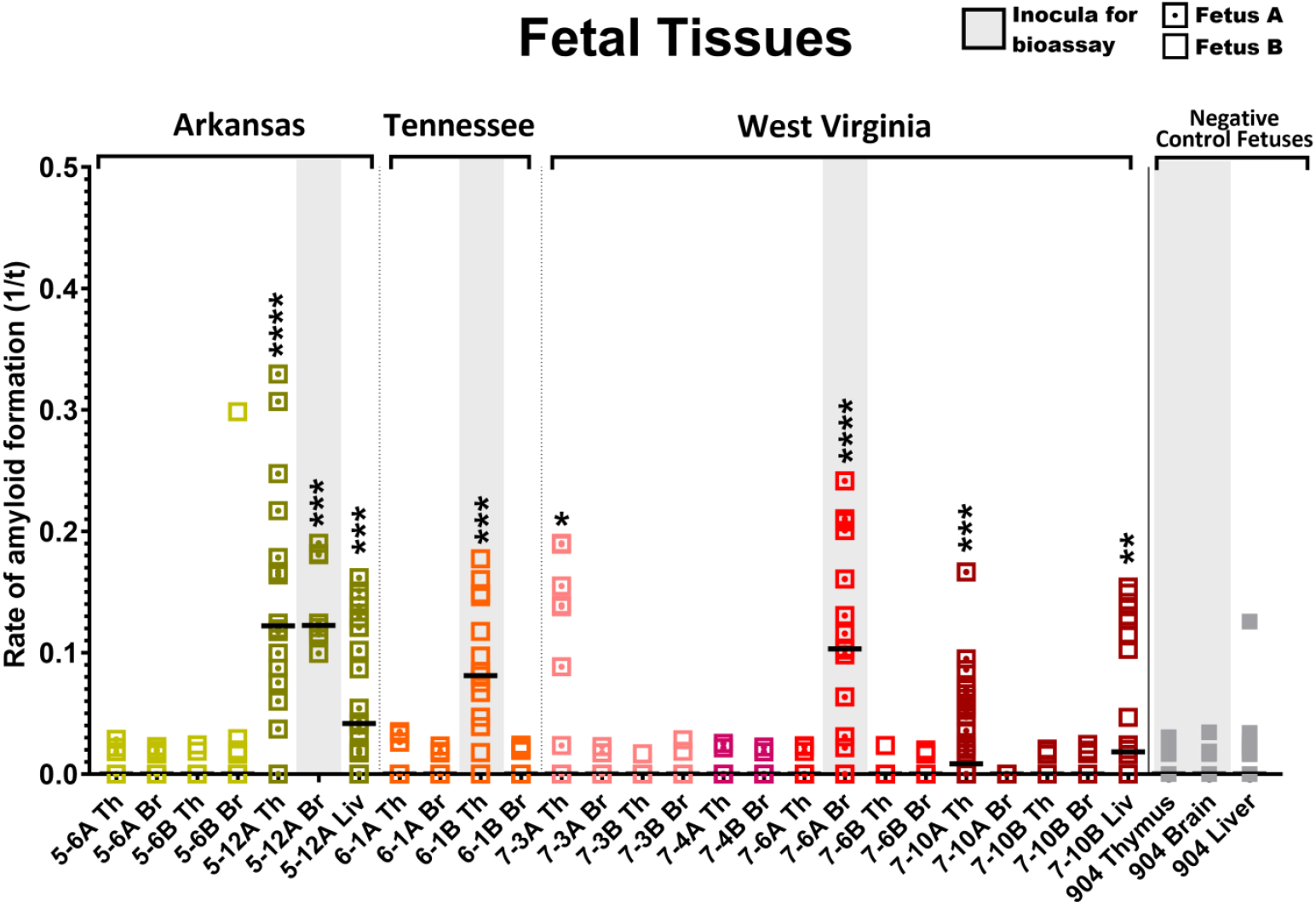
CWD detection in white-tailed deer (WTD) fetal tissues determined by RT-QuIC. Amyloid seeding detected in fetal brain, thymus, and liver (Table 2) collected from CWD-positive dams as determined by RT-QuIC and IHC (Table 1). Amyloid formation displayed as reaction rates (1/time (t) to ThT fluorescent threshold). Horizontal lines represent medians with n=16 replicates (fetal tissues). Negative control fetal tissue samples from WTD collected in Georgia (n=3) remained negative in all samples tested. P values as follows: * <0.05, ** <0.01, *** <0.001, ****<0.0001. Grey shaded rectangles denote samples utilized for bioassay inoculations (Table 4). Abbreviations represented as follows: (Th)ymus, (Br)ain, and (Liv)er.

### Infectious prions present within the pregnancy microenvironment and fetuses harvested from free-ranging white-tailed deer

To determine the biological relevance of the presence of *in vitro* conversion competent prions in maternal and fetal tissues, we assessed a subset of tissues harvested from the reproductive tract (uterus), the maternal-fetal interface (placentome), and fetuses (thymus and brain) for the presence of the infectious CWD prions in Tg^CerPrP-E226^5037^+/-^ mouse bioassay.

Pregnancy microenvironment: All mice (35/35; 100%) inoculated with uterine or placentome tissue developed terminal clinical TSE disease (ataxia, limb paralysis, weight loss, etc.); uterine tissue (12/12; 264 ± 30 days post-inoculation (dpi) (mean ± SD)), placentome tissue (23/23; 256 ± 27 dpi). *In vitro* amyloid seeding activity was present in brain tissues harvested from all mice (35/35) as assessed by RT-QuIC (**Table 3**).

Fetal tissues: Six of seven (6/7; 85.7%) mice inoculated with fetal thymus (325 ± 44 dpi) and 7 of 11 (63.6%) inoculated with fetal brain (254± 80 dpi) developed terminal clinical disease which was confirmed in brain and/or spleen by RT-QuIC (**Table 4**). Age-matched and mock-inoculated negative control mice remained free of clinical signs of disease throughout the 381 dpi study and were confirmed negative by RT-QuIC (**Figure 3)**.

**Table 4.**
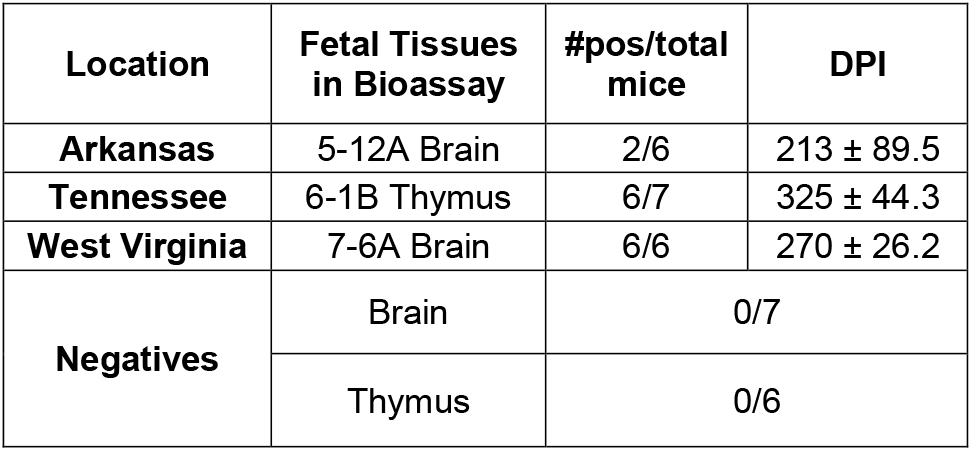
Mouse bioassay results for white-tailed deer fetal tissues from CWD-positive dams. Cervidized (Tg^CerPrP-E226^5037^+/-^) mice were intraperitoneally (IP) inoculated with fetal brain (fetuses 5-12A and 7-6A) or fetal thymus (6-1B) samples. Terminal mouse brain and spleen tissues were tested in conventional RT-QuIC for amyloid seeding activity. Mock inoculated mice remained free of clinical signs throughout, and tested CWD-negative upon terminal collections in all cohorts. DPI = days post inoculation. # pos/total mice = number CWD-positive/total number inoculated.

| Location | Fetal Tissues in Bioassay | #pos/total mice | DPI |
| --- | --- | --- | --- |
| Arkansas | 5-12A Brain | 2/6 | 213 ± 89.5 |
| Tennessee | 6-1B Thymus | 6/7 | 325 ± 44.3 |
| West Virginia | 7-6A Brain | 6/6 | 270 ± 26.2 |
| Negatives | Brain | 0/7 |  |
|  | Thymus | 0/6 |  |

**Figure 3.**
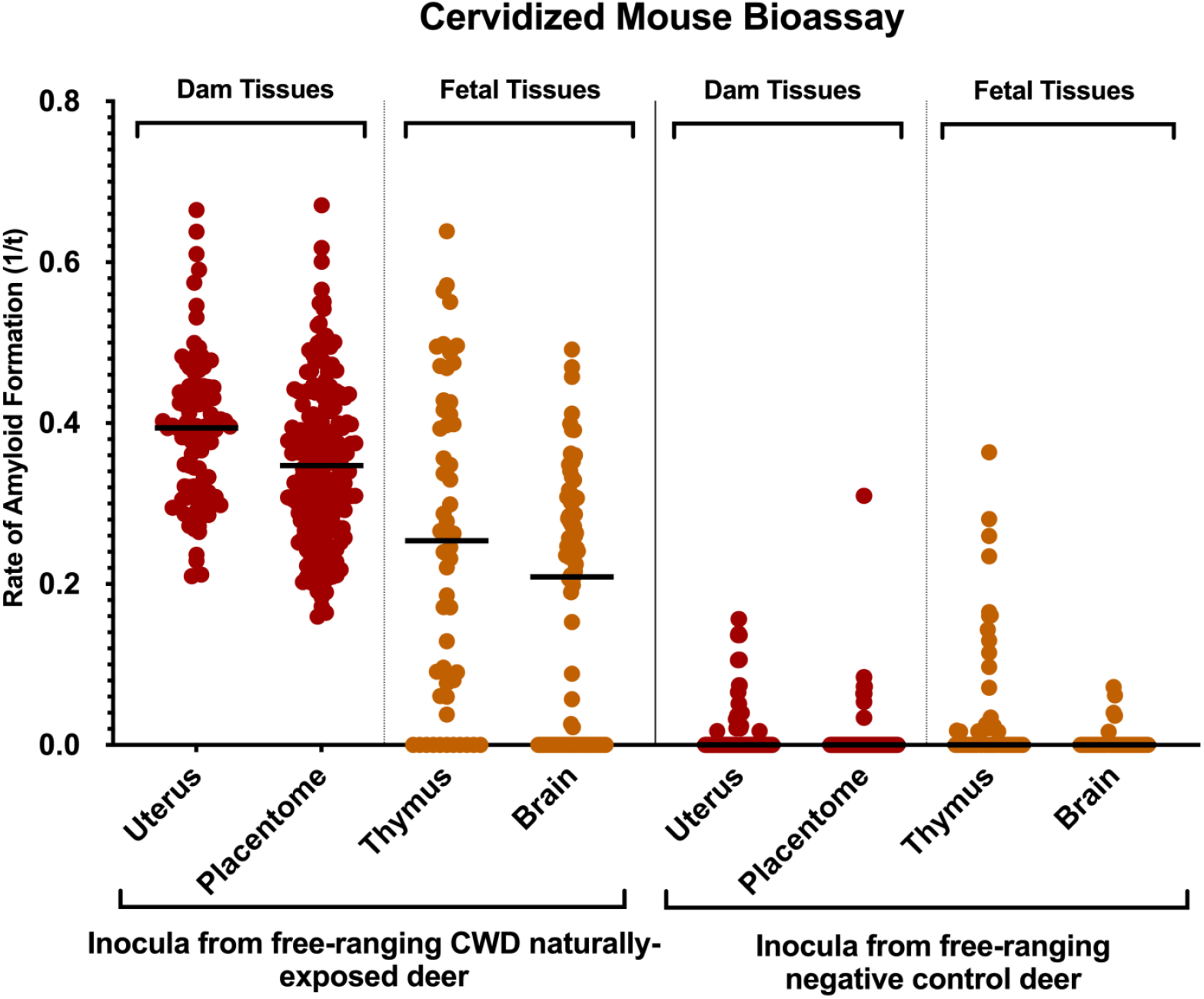
Infectious CWD prions present in white-tailed deer maternal reproductive and fetal tissues determined by mouse bioassay. Cervidized (Tg^CerPrP-E226^5037^+/-^) mice were inoculated with 10^−4^ dilutions of uterus or placentome intracranially (uterus: n=12 mice; placentome: n=35 mice) or fetal thymus or fetal brain intraperitoneally (thymus: n=7 mice; brain: n=11 mice). Terminal mouse brains were tested by conventional RT-QuIC for amyloid seeding activity. Amyloid formations as reaction rates (1/time (t) to ThT fluorescent threshold). Horizontal lines represent the medians (8 replicates per mouse). Mice inoculated with tissues from free-ranging negative control deer (uterus: n=8 mice; placentome: n=8; thymus: n=7; brain: n=6; 8 replicates each) from Georgia served as negative controls. A Mann-Whitney unpaired t-test was used against negative controls to determine statistical significance. **** indicates a P value of <0.0001.

In summary, we demonstrate that free-ranging WTD naturally exposed to CWD harbor infectious CWD prions in maternal and gestational fetal tissues.

### Gestational Age

Gestational age, as determined by the Hamilton^35^ equation of rump to head measurements, showed fetal age ranging between 79 to 158 days from conception, with an average age of 112.5 days (total gestational time for WTD is 201 days). There were no fetuses harvested in the first trimester of development (0-67 days), n=44 fetuses harvested in the second trimester (68-134 days), and n=10 fetuses harvested in the third trimester (135-201 days). The age-match negative control Georgia fetuses (n=3) were an average age of 103.5 days. We found tissue positivity in fetuses at gestational age 90 to 158 days, correlating to 2^nd^ and 3^rd^ trimester pregnancies.

### *Prnp* genotypes from free-ranging white-tailed deer dams and fetuses

As *Prnp* genotypes have been shown to be important in host susceptibility to prion infection and *in utero* transmission of scrapie in sheep, we evaluated maternal (n=31) and fetal (n=54) tissues to determine their genotypes at codon 95 and 96. The *Prnp* open reading frame was sequenced for all the WTD to determine the impact genotype (codon 95/96) may have on vertical transmission. Pregnant dams in this study (n=28 CWD endemic; n=3 nonendemic) represented all three polymorphisms at codon 96: GG=21, GS=9, and SS=1. When *Prnp* polymorphisms were correlated with dam CWD positivity, we identified CWD-positive dam status in 12 of 21 (57.1%) codon 96GG, 4 of 9 (44.4%) codon 96GS, and 0 of 1 (0%) codon 96SS (**Table 1**). All dams were wildtype QQ at codon 95 (31/31).

To determine if polymorphisms at *Prnp* codons 95 and 96 may play a role in CWD transmission and deposition in the fetus, we determined the *Prnp* genotype of each fetus in this study. Of the 54 fetuses collected, all three genotypes were present at codon 96; GG=36, GS=16, and SS=2. When compared to their CWD status, all 6 CWD positive fetuses were Prnp codon 96GG (6/36;16.7%), with no CWD detection in fetuses expressing 96GS or 96SS polymorphisms (**Table 2**). Interestingly, there were two instances of a mutation in the *Prnp* 95 codon where Glutamine was substituted for Histidine (QH). This was the case in one CWD positive fetus from a CWD positive dam (**Table 2**) and in one CWD negative fetal/dam dyad. The remaining fetuses were wildtype homozygous QQ at the same position (n=52). Our study suggests that vertical transmission may be affected by variations at codon 96, as there were no cases of fetal positivity in dams with *Prnp* genotypes codon 96 GS and SS; polymorphisms that convey delayed disease progression.

### Historical CWD surveillance data

All three states in the study area have historically conducted limited CWD ELISA and/or IHC testing on fawns as part of state wildlife agency CWD surveillance and monitoring programs (**Table 5**). CWD was detected in lymphoid (retropharyngeal lymph node; AR, TN, WV) or brain (obex; WV) tissues harvested from fawns (total 74/1476; 5%) that varied in age from <6 to 10 months.

**Table 5.** Historical chronic wasting disease (CWD) test results for fawns (≤ 10 months) from each of the three study areas in Arkansas, Tennessee, and West Virginia. The data were collated from wildlife management agency surveillance efforts over varying amounts of time. Retropharyngeal lymph nodes or obex were tested by either enzyme-linked immunosorbent assay (ELISA) and/or immunohistochemistry (IHC) depending on agency protocol.

| Years | State | County | Age | Tests | No. Positive | No. Tested |
| --- | --- | --- | --- | --- | --- | --- |
| 2016-2024 | Arkansas | Newton | ≤6 mo | ELISA | 27 | 231 |
| 2018-2024 | Tennessee | Fayette | 5-8 mo | ELISA | 32 | 966 |
| 2006-2012, 2017, 2022 | West Virginia | Hampshire | 8-10 mo | ELISA and/or IHC | 15 | 279 |

In summary, we demonstrate *in vitro* amyloid seeding in the pregnancy microenvironment of half of the confirmed CWD-positive free-ranging dams tested, with a third of these dams carrying at least one fetus with PMCA amplifiable prion seeds. Of most importance, a subset of fetal tissues inoculated into bioassay confirms the presence of infectious CWD prions in maternal and gestational fetal tissues providing the first evidence of the initiation of CWD infections in the fetus prior to parturition. To add additional biological relevance to these findings, state wildlife agency CWD surveillance and monitoring programs have detected CWD in fawns ≤ 10 months of age.

## DISCUSSION

The continued detection of CWD in free-ranging cervid populations and increasing prevalence in some locations demonstrate the need for a greater understanding of the impact maternal infections have on the transmission of this fatal neurodegenerative disease. With CWD prevalence as high as 50% in some localized areas of the U.S.^39^ understanding how mother-to-offspring transmission impacts the overall transmission dynamics of this disease will be imperative to those who enjoy, hunt, oversee, and manage captive and wild cervid populations. To this end, this multi-state and -agency study came together to determine if healthy-appearing female white-tailed deer from CWD endemic regions, that are naturally exposed to CWD, can transmit the disease to their fetus during pregnancy.

Here we report the presence of the infectious CWD agent within fetal and reproductive tissues of free-ranging WTD, revealing that dams can transmit the disease *in utero* to their offspring. In agreement with previous studies^30,31^ we also confirm CWD deposition within reproductive maternal tissues (**Figure 1**) and *in utero*-derived fetal tissues (**Figure 2**) harvested from free-ranging cervids using highly sensitive *in vitro* amyloid detection assays. Although this and previous studies have shown PrP^CWD^ deposition in tissue, what was left unknown, was whether these detections represented infectious prions capable of initiating and progressing CWD infections. We employed mouse bioassays, regarded as the only tool to determine the presence of prion infectivity, and found that female reproductive (i.e., uterus, placentome, amniotic fluid) and fetal tissues (i.e., thymus, brain, liver) harbor infectious CWD prions (**Figure 3, Tables 3, 4**). Thus, this study provides strong support that offspring can be exposed to and initiate CWD infections long before parturition.

Interestingly, our study reveals a nuanced picture of CWD transmission in the pregnancy microenvironment of cervid placentomes. We observed variable positivity within placentomes of fetal twin sets, as well as within each fetus’ own placentome structure, confirming findings from previous studies in muntjac and rocky mountain elk^28,29,32^. In addition, we demonstrate that prion deposition within the reproductive microenvironment (i.e., placentomes, uterus and amniotic fluid) does not always result in prion deposition within the fetus. This suggests other factors, such as maternal and paternal genetic input, may contribute to the susceptibility of vertical transmission. In these cases it is possible that CWD progression within this environment has been slow to progress, or as noted in this study, may be associated with maternal-fetal dyads that express *Prnp* 96GS haploid, a polymorphism in the cervid prion protein known to delay disease onset^33,34^. The observation could hint at difficulties in overcoming the protective influence of 96GS genotypes in maternal transmission. Yet, other studies have shown evidence that GS 96 fetuses contain CWD prions^30^, indicating that, similar to adult WTD, protective effects of the polymorphism are not enough to prevent *in utero* transmission of the disease. We also demonstrate a CWD-positive fetus with the protective 95QH polymorphism, indicating that *in utero* transmission can also overcome this genetic variant (**Table 2**). These findings are in agreement with those conducted in the related sheep scrapie system^26^, where it was demonstrated that sheep with *Prnp* polymorphisms that confer protection from disease resulted in reduced or absent *in utero* transmission to lambs. Also, gestational age did not have a correlative effect on fetal CWD status, showing that fetuses can be infected during the 2^nd^ and 3^rd^ trimesters of gestation (**Table 2**). We have previously reported this in experimental muntjac studies demonstrating CWD deposition in *in utero*-derived fetuses harvested from CWD-positive dams throughout all periods of gestation (1^st^, 2^nd^ and 3^rd^ trimester)^28^, while studies in the sheep scrapie system^26^ report progressively increasing placentome PrP^Sc^ deposition from early to term pregnancy.

Horizontal transmission has long been recognized as the primary driving force in the spread of CWD, yet vertical transmission continues to gain traction as a contributing mechanism. Vertical transmission has been shown to be an efficient route of transmission in several prion diseases including scrapie in sheep^20,21,40-44^, experimental CWD in Reeves’ muntjac^29^, and has been suggested for CWD in captive and free-ranging WTD^30,31^ and free-ranging Rocky Mountain elk^32^. The results reported here, evaluating free-ranging animals, are consistent with previous experimental cervid studies revealing the presence of prion seeding and infectious CWD agent within maternal uterus, placentomes, and amniotic fluid^29^. These findings will further our understanding of CWD prion transmission at a population level and enable investigation of the potential for influence on population dynamics. Experimental studies conducted in the Reeves’ muntjac demonstrate that muntjac at all stages of CWD infection were able to breed and maintain full term pregnancies^28^. However, these studies also revealed a 60% increase in nonviable offspring born to CWD-infected muntjac dams. If this is representative of white-tailed deer and other cervid species in a free-ranging setting, it may suggest a negative impact to recruitment. This will be an important, but difficult area of study. In mule deer and WTD, subtle negative effects on fawn survival and recruitment have been reported^45^. As CWD is known to result in population declines in mule deer and white-tailed deer^46,47^, there is need to understand if fetal infections contribute to such declines.

Questions remain about the impact of these findings on free-ranging cervid populations. Here, pregnant WTD were harvested from three southeastern U.S. states (Arkansas, Tennessee, and West Virginia) in areas with apparent CWD prevalence exceeding 20% to assess impacts of maternal infections. Our findings provide evidence that maternal CWD infections within these states are resulting in gestational infections demonstrating prion exposure and amplification within fetal tissues long before parturition. Our previous experimental studies in muntjac revealed that offspring born to CWD-infected dams became lymphoid tissue positive as early as 41 days after birth, and progressed to terminal clinical disease between 2-5 years of age^28^. Information regarding progression of *in utero* infection to a disease state after parturition is not known for free-ranging WTD populations; however, this is an important area of inquiry. Prion transmission from healthy-appearing CWD-infected dams to their offspring has the potential to influence future fawn recruitment and prevalence rates in conjunction with infections acquired via horizontal means. If the timing and progression of clinical CWD is such that offspring die from CWD earlier than they would from an indirect exposure and prior to replacing themselves in the population, recruitment can be impacted through decreased lifetime reproductive potential in females.

Our detection of CWD-positive fawns ≤10-month-old by ELISA testing provides indirect evidence of gestational infection, although direct or indirect CWD transmission after birth cannot be ruled out in these fawns (**Table 5**). The detection of CWD infection in ≤6-month-old fawns in the Arkansas and Tennessee study sites by ELISA test, which is not as sensitive as amplification assays, was particularly suggestive of vertical transmission. Fawns less than 6 months of age are not routinely incorporated into statewide CWD surveillance programs because of the low likelihood of detectable infection using traditional assays (i.e., ELISA, IHC), yet have been previously identified^17^. Future studies should investigate the potential role of *in utero* transmission in local CWD transmission cycles. Considering the strong relationship between CWD infection probability and female white-tailed deer relatedness, *in utero* CWD transmission may serve as an important route of transmission in addition to social interactions and indirect exposures^48^.

Overall, this study describes the dissemination of CWD prions throughout tissues and birthing fluids of the pregnancy microenvironment demonstrating that offspring are routinely exposed to the infectious prion *in-utero* prior to parturition. We report infectious prions in the reproductive and fetal tissue of naturally exposed free-ranging white-tailed deer suggesting that *in utero* maternal transmission is likely an underappreciated mode of CWD transmission. Our study shows that vertical transmission is indeed a viable route of infection within the southeastern U.S. and is another factor contributing to the relentless spread of chronic wasting disease.

## METHODS

Research reported in this publication was supported by the Wildlife and Sport Fish Restoration Programs of the U.S. Fish and Wildlife Service (Award number F20AP00172) and by the National Institute of Allergy and Infectious Diseases of the National Institutes of Health under Award Numbers R01AI112956, P01-AI-077774, and R01AI156037. The content is solely the responsibility of the authors and does not necessarily represent the official views of the National Institutes of Health. Additional support was provided by SCWDS member states (Alabama, Arkansas, Florida, Georgia, Kentucky, Kansas, Louisiana, Maryland, Mississippi, Missouri, Nebraska, North Carolina, Oklahoma, South Carolina, Tennessee, Virginia, and West Virginia), U.S. Fish and Wildlife Service National Wildlife Refuge System, and U.S. Geological Survey Ecosystems Mission Area. Field sampling was approved by the University of Georgia’s IACUC # A2018 02-010. Mouse bioassay was approved by Colorado State University’s IACUC #1524 and 5362. The authors complied with the ARRIVE guidelines.

### Study area, sample collection & preparation

Three CWD endemic areas of the southeastern United States were targeted for lethal collection of free-ranging, adult, female WTD, including: Newton County, Arkansas; Fayette County, Tennessee; and Hampshire County, West Virginia. Mature females were targeted January through April in these areas because the apparent CWD prevalence in WTD at each collection site was greater than 20%, optimizing chances of collecting CWD-positive gravid WTD between 16-24 weeks gestation. Deer were similarly collected from a control site in Clarke County, Georgia, a location with no detection of CWD in WTD. All deer were lethally collected by state wildlife management agency or Southeastern Cooperative Wildlife Disease Study (SCWDS; University of Georgia) personnel. Collection and sampling occurred across a calendar year, including January 2020 in Georgia (n=3 dams; n=3 fetuses), February 2020 in Arkansas (n=10 dams; n=17 fetuses), March 2020 in Tennessee (n=10 dams; n=20 fetuses), and April 2021 in West Virginia (n=8 dams; n=14 fetuses).

Adult WTD necropsies and sample collection were performed by SCWDS personnel at each field site. Care was taken to use single-use instruments, single-use personal protective equipment, disposable supplies, intentional workflow, and proper technique to minimize risk of cross-contamination between animals, as well as between tissues within individual animals. Immediately upon death, whole blood was collected by cardiocentesis using a needle and syringe. The blood was transferred to vials containing citrate for processing and CWD status testing by prion amplification assays. Saliva (1-2 mL) was collected using disposable transfer pipettes from the buccal cavity and transferred to cryovials for CWD testing by prion amplification assays. Each carcass was placed into a body bag and transported by vehicle to a field necropsy and sample collection site. Age of each animal was estimated based on tooth wear and replacement^49^. A small team of individuals with assigned roles sampled deer individually and collected brainstem, retropharyngeal lymph nodes (RPLN), palatine tonsils, rectoanal mucosa-associated lymphoid tissue (RAMALT) and third eyelids. Tissues were split between fresh-frozen and 10% neutral buffered formalin for later detection of PrP^CWD^ by immunohistochemistry (IHC) or prion amplification assays (RT-QuIC, PMCA). Two plastic zipties were secured to the cervix and the intact, gravid uterus was excised and placed into large plastic bags. Urine (1-2 mL) was collected from the urinary bladder using needle and syringe and transferred to a cryovial. Blood, saliva, urine, intact uterus, and all maternal tissue samples were stored at -20°C overnight prior to shipment to Colorado State University (CSU) where they were stored at -80°C until further processing.

At CSU, tissue samples were thawed and intact uteri and fetuses were individually necropsied by CSU personnel using single-use tissue and fetus instruments to mitigate cross-contamination between maternal and fetal tissues. Fetuses were placed at 4°C overnight to thaw. Amniotic fluid was extracted by use of single-use needle and syringe from each amniotic sac. Each amniotic sac was opened using a single-use scalpel and fetuses were removed and measured for gestational age using the Hamilton equation^35^; (Age [days]= (body length [mm] x 0.32) + 36.82). Fetuses were carefully dissected using new PPE and sterile single-use instruments between each fetus and their respective tissues; all collected items were stored at -80°C for future analysis. All tissues were assessed by the prion amplification assays protein misfolding amplification assay (PMCA) and/or real-time quaking induced conversion (RT-QuIC) for the presence of amyloid seeding activity. Dam and fetal tissue were made into 10% w/v tissue homogenates with 1x PBS (20 mM NaPO_4_, 150 mM NaCl, pH 7.4) in an Omni Bead Ruptor 24 and stored at -80°C until tested. The following tissues and fluids were analyzed from pregnant dam: obex, RAMALT, RPLN, tonsil, third eyelid, blood, uterus, placentomes, amniotic fluid, saliva, and urine. The following tissues were analyzed from each fetus: brain, thymus, liver, and spleen. Independent researchers assessed maternal and fetal tissues and remained blinded to outcomes until the study was completed.

### Immunohistochemistry (IHC)

IHC is the gold standard for PrP^CWD^ detection and was used to visualize prion deposition within fixed tissue. Paraffin embedded WTD dam tissue sections of RPLN, RAMALT, obex, third eyelid, or tonsil were deparaffinized with xylene (100%) and rehydrated in graded alcohols (100% - 70%) before treatment with 88% formic acid (30 minutes at room temperature) as previously described^50^. After washing with water, slides were placed in a 2100-Retriever (Prestige Medical) in sodium citrate buffer (0.01M sodium citrate, 0.05% Tween 20, pH 6.0) for heat-induced epitope antigen retrieval. Endogenous peroxidase activity was quenched with 3% hydrogen peroxide and slides were blocked before overnight incubation in anti-prion antibody BAR224 (1 mg/mL; Caymen Chemical) diluted 1:750. Envision+ anti-mouse HRP-labeled polymer secondary antibody (Dako) was followed by 3-amino-9-ethylcarbazole (AEC) (Dako) for visualization. Negative control tissues were run concurrently with test samples.

### Iron Oxide Bead Extraction (IOB)

As prion burdens are known to be relatively low in bodily fluids, we previously developed a pre-RT-QuIC concentration method, iron oxide bead (IOB) extraction, to enhance detection capability^51^. Amniotic fluid diluted 1:10 and 1:100 (1x PBS), saliva diluted 1:20 (1x PBS), or undiluted urine (1 mL total volume/each), were added to individual 1.7 ml conical tubes^36,51^. Two microliters (2 µL) of iron oxide beads (BM547 Bangs Laboratory) were added to each tube followed by end-over-end mixing at room temperature for 30-60 minutes. Tubes containing samples were placed on a magnetic particle separator (Pure Biotech, New Jersey) which permitted the bead bound prions to be immobilized against the tube wall. Supernatants were discarded and the tubes were removed from the magnet. The beads, containing extracted prions, were resuspended in 10 µL of 0.1% sodium dodecyl sulphate (SDS) and run in RT-QuIC at 42°C. Positive and negative controls were run with all IOB-QuIC plates.

### Real-Time Quaking Induced Conversion (RT-QuIC)

We used RT-QuIC to determine CWD status of fetal and maternal tissues, as previously described^29,52^. Briefly, 0.1 mg/mL of truncated Syrian hamster recombinant protein containing amino acids 90-231 was added to master mix (20 mM NaPO_4_, 1 mM ethylenediaminetetraacetic acid tetrasodium salt (EDTA) (Sigma), 320 mM NaCl, and 10 µM thioflavin T (ThT)). Master mix (98 ul) was added to each well of a black optical bottom 96-well plate. Two microliters (2 µl) of maternal tissue homogenates at a final dilution of 10^−3^ (third eyelid, uterus, placentomes), 10^−4^ (RAMALT, RPLN, tonsil), or 10^−5^ (obex), while 2 µl fetal tissues prepared from round five PMCA product (final dilution of 10^−2^ in 0.1% SDS in PBS) were loaded into each well in quadruplicate. Reproductive and fetal tissues were tested by two independent researchers blinded to the results of the dam tissues. Plates were inserted into a BMG Fluostar Omega plate reader and analyzed at 15-minute intervals for 250 rounds or 62.5 hours at 42°C or 55°C as previously optimized for each tissue type and protocol^51,53^. A minimum of two independent quadruplicate plates were run per tissue sample for a total of 8 replicates per tissue. CWD positive and negative WTD tissue specific controls were assessed with all maternal tissues. Reeves’ muntjac and/or WTD specific fetal tissues served as positive and negative controls with all fetal tissues. Reactions were considered positive when fluorescent readouts reached five standard deviations above the average initial five fluorescent readings. Significance was determined by comparing age-matched negative control tissues to samples of interest using the unpaired, non-parametric, two-tailed Mann-Whitney Test in GraphPad Prism v9 to generate *p*-values. Significance was represented as follows: \**P*≤0.05, \*\**P*≤0.01, \*\*\**P*≤0.001, \*\*\*\**P*≤0.0001.

### Protein Misfolding Cyclic Amplification (PMCA)

To further investigate prion burden in fetal tissues we subjected each to an additional amplification method, PMCA. Normal brain homogenate (NBH) was prepared from naive transgenic mice that over-expressed cervid PrP (Tg^CerPrP-E226^5037^+/-^) to serve as PMCA conversion substrate. Briefly, mice were perfused with 5 mM EDTA prior to whole brain extraction. Brain tissue was flash frozen in liquid nitrogen, and subsequently made into a 10% w/v homogenate (5mM EDTA, 150 mM NaCl and 1% Triton X-100). Fetal tissue homogenates (10% w/v) and NBH were added in 0.2 mL microcentrifuge tubes containing 2.38 mm and 3.15 mm Teflon beads (McMaster-Carr)^37,54^. The first round of PMCA was initiated with the addition of fetal tissue homogenates at a ratio of 10 µL tissue sample to 90 µL NBH (exception: 10 µL spleen tissue diluted to 10^−2^). PMCA rounds two through five were comprised of 20 µL previous rounds’ amplified product to 50 µL fresh NBH substrate^37^. Round one PMCA samples were placed in a Misonix sonicator for 72 hours (144 cycles; each cycle consisting of 30 seconds sonication and 29.5 minutes incubation at 37°C). PMCA rounds two through five were processed for 24 hours (48 cycles; each cycle consisting of 30 seconds sonication and 29.5 minutes incubation at 37°C). All samples were run with CWD positive and negative tissue specific controls in duplicate by two independent investigators.

### sPMCA with RT-QuIC (PQ)

We have previously demonstrated enhanced prion detection sensitivity by combining PMCA with RT-QuIC ^55,56^. Here, round five PMCA product was further prepared for amyloid seeding assessment by RT-QuIC with each sample being diluted 1:100 in 0.1% SDS. Each sample was prepared and assessed as per the RT-QuIC methods above.

### Bioassay in cervid transgenic mice

Bioassay in natural host or rodent model species is the only way to determine the presence of the infectious prion agent has the capacity to initiate and progress disease. Thus, we used mouse bioassay to determine the presence of the infectious CWD agent within tissues of the pregnancy microenvironment and fetus. Uterine and placentome tissues (pregnancy microenvironment) harvested from CWD naturally infected free-ranging dams were intracranially (IC)-inoculated into Tg^CerPrP-E226^5037^+/-^ mice. Mice were inoculated at six to twelve weeks of age and maintained as previously described^29^ and approved by CSU’s IACUC. Each mouse was IC-inoculated in the right parietal lobe with 30 µl of 10% w/v homogenate of uterus or placentome (n=12, n=23 mice, respectively) or intraperitoneal (IP)-inoculated with fetal brain or fetal thymus (n=11, n=7 mice, respectively) using 80 µL of 1:2.5 round 5 PMCA material diluted in PBS that was treated overnight with 2% HyClone Pen-Strep. All mice were monitored daily for signs of prion infection (ataxia, weight loss, hypersensitivity) until termination due to clinical disease presentation. Dam pregnancy microenvironment tissues (uterine and placentome) and *in utero*-derived fetal tissues harvested from white-tailed deer in Clarke County, Georgia (n=3) were treated identically and either IC or IP-inoculated into cohorts of mice (n=16, n=13 mice, respectively) serving as negative bioassay controls. Brain and spleen tissue harvested from each mouse at termination were assessed for amyloid seeding (prions) by RT-QuIC.

### Genotyping

As polymorphisms within the *Prnp* coding sequence are known to affect disease characteristics^34,57^, all dams and fetuses were genotyped with codon 95 and 96 reported for this study. DNA was extracted from dam (RPLN or spleen) and fetal (thymus or spleen) tissues using a DNeasy Blood and Tissue kit (QIAGEN). DNA concentration was determined for each sample (NanoDrop – 1000 UV/Vis Spectrophotometer) prior to polymerase chain reaction (PCR) to amplify the purified DNA. Deer specific PCR primer pair 213d *AGGTCAACTTTGTCCTTGGAGGAG* and 139u *TAAGCGCCAAGGGTATTAGCAT*, with sequencing primer 86d *CAGTCATTCATTATGCTGCAGACT* were used as previously described^28^. PCR amplified DNA was sent for Sanger sequencing by Quintara Biosciences and the returned results were analyzed using the sequencing software Chromas (Technelysium Pty Ltd v2.6.6).

### Historical CWD surveillance data

Available CWD surveillance data from the three study areas was accessed via each state wildlife agency database. Data sources varied for each state. For Arkansas, county-level data for Newton County were examined, including all available sources for WTD ≤ 6 months (e.g., hunter-harvest, road-kill, sick/target deer). Samples in Arkansas were tested by ELISA and/or IHC. In Tennessee, data from hunter-harvested fawns from Fayette and Hardeman counties were assembled. Tennessee fawns were tested by ELISA. In West Virginia, CWD data was assembled from routine agency special collections in a 39 mile square area of Hampshire County. All deer were <6-10 months old at the time of collection and were tested by ELISA and/or IHC. All assays were performed routinely at accredited veterinary diagnostic laboratories.

## DATA AVAILABILITY

The datasets generated during and/or analyzed during the current study are available from the corresponding author on reasonable request.

## ACKNOWLEDGEMENTS

Funding for this research was provided by the Multistate Conservation Grant Program through the Wildlife and Sport Fish Restoration Programs of the U.S. Fish and Wildlife Service (grant # F20AP00172). Additional support was provided by the long-term financial support of SCWDS by member agencies in Alabama, Arkansas, Florida, Georgia, Kansas, Kentucky, Louisiana, Maryland, Mississippi, Missouri, Nebraska, North Carolina, Oklahoma, South Carolina, Tennessee, U.S. Virgin Islands, Virginia, and West Virginia coordinated under the Federal Aid in Wildlife Restoration Act (50 Stat. 917), and the U.S. Geological Survey Ecosystems Mission Area and U.S. Fish and Wildlife Service National Wildlife Refuge System. Further financial support was provided by the National Institute of Allergy and Infectious Diseases of the National Institutes of Health under Award Numbers R01AI112956, P01-AI-077774, and R01AI156037. The content is solely the responsibility of the authors and does not necessarily represent the official views of the National Institutes of Health. The views and opinions expressed herein are those of the authors and do not necessarily reflect the views or policies of the Arkansas Game and Fish Commission. Product references do not constitute endorsements. We thank Michael Chamberlain for continued support of this work and the staff for their assistance in the field, especially John Wlodkowski (SCWDS); TWRA CWD field staff; Wes Wright and Stacey Clark (AGFC); Rich Rogers, Lee Strawn, and District 2 staff (WVDNR). We thank Joseph Westrich for critical review of this manuscript.

## AUTHOR CONTRIBUTIONS

AMS: data acquisition, analysis, interpretation, and manuscript drafting, AVN: data acquisition, analysis, interpretation and manuscript drafting, EEM: data acquisition, analysis, interpretation and manuscript review, NDD: data acquisition, analysis, interpretation and manuscript review, DJT: data acquisition, interpretation, and manuscript review, ZO: data acquisition, EB: animal collection, data acquisition and manuscript review JB: data acquisition and manuscript review DMG: design, facilitated field collections, manuscript review JSD: animal collection, data acquisition, manuscript review NS: design, data acquisition, manuscript review CAC: design, data acquisition, manuscript review JMC: state agency coordination, animal collection, financial grant collaborator, manuscript review MGR: study conception, financial support, data acquisition, analysis, interpretation, manuscript drafting, CKM: study conception, design, data analysis and interpretation, manuscript drafting, and financial support.

